# Myeloid cell mPGES-1 mediates inflammatory pain hypersensitivity in mice

**DOI:** 10.1101/2020.01.31.929422

**Authors:** Lihong Chen, Guangrui Yang, David P. Cormode, Ashmita Saigal, Shravanthi Madhavan, Liudmila L. Mazaleuskaya, Gregory R. Grant, Garret A. FitzGerald, Tilo Grosser

## Abstract

Nonsteroidal anti-inflammatory drugs (NSAIDs) relieve inflammatory pain by predominant suppression of cyclooxygenase-2 derived prostaglandin (PG) E_2_. Innate immune cells contribute to inflammatory pain hypersensitivity and may be an attractive target for novel non-addictive approaches to pain management. We studied the contribution of PGE_2_ produced by myeloid cell microsomal prostaglandin E synthase -1 (mPGES-1) to peripheral inflammation and hyperalgesia in mice. Selective deletion of mPGES-1 in myeloid cells by crossing LysM-Cre mice with mPGES-1flox/flox mice (Mac-mPGES-1-KO) resulted in significantly reduced mechanical and thermal hyperalgesia in complete Freund’s adjuvant (CFA)-evoked hind paw inflammation, zymosan-induced peri-articular inflammation and collagen II antibody-induced arthritis models. Systemic COX-2 inhibition or myeloid cell specific COX-2 deletion (by crossing LysM-Cre with COX-2 flox/flox mice) recapitulated reduction of CFA-induced inflammation and hyperalgesia. In contrast, deletion of mPGES-1 in neurons and glial cells by crossing mPGES-1flox/flox mice with Nestin-Cre mice had no detectable effect on inflammatory pain hypersensitivity. While macrophage recruitment was unaltered, tissue concentrations of PGE_2_, IL-1β and TNFα were significantly reduced in Mac-mPGES-1-KO paw tissues following CFA induction. Our results demonstrate that myeloid cell mPGES-1 is the dominant source of PGE_2_ in inflammatory pain hypersensitivity. Targeting myeloid cell mPGES-1 may afford a novel approach to inflammatory pain therapy.

## Introduction

Nonsteroidal anti-inflammatory drugs (NSAIDs) are on average as effective, or more so, than immediate-release opioids for pain that is primarily driven by inflammatory processes (1,2). The analgesic efficacy of NSAIDs has been demonstrated in headache including migraine, dysmenorrhea, traumatic and post-operative pain, rheumatoid arthritis, osteoarthritis and ankylosing spondylitis (3), although there is considerable interinvidual variability in pain relief (4). NSAIDs inhibit the formation of cyclooxygenase-(COX) derived prostanoids involved in the sensitization of the nociceptive system (5). As immune surveillance cells recognize tissue injury, they initiates an inflammatory response of the innate immune system to remove cell debris and begin the healing process. This process is highly regulated and involves the recruitment of the COX biosynthetic-response network, in which COX-2 in particular is induced in cytokine-activated tissue resident and infiltrating immune cells, including macrophages, neutrophils and mast cells (6). COX products, most notably prostaglandin (PG) E_2_ lower the activation threshold of ion channels involved in sensing pain and increase the gain of the pain signaling system (5).

Microsomal PGE Synthase-1 (mPGES-1) is downstream of the COXs and the dominant source of PGE_2_ biosynthesis in inflammation in comparison to mPGES-2 and cytosolic (c) PGES (7). Due to its preferential coupling with COX-2, mPGES-1 has received much attention as a potential drug target for novel analgesic and anti-inflammatory drugs (7). Global deletion of mPGES-1 has been as effective as NSAID treatment in rodent models of pain and inflammation (8,9) and specific inhibitors of this enzyme have been synthesized and advanced into clinical development (10–13). The expectation has been that inhibition of mPGES-1 would retain the analgesic efficacy of NSAIDs, but reduce the likelihood of gastrointestinal and cardiovascular adverse effects (7).

Indeed, deletion of mPGES-1 in endothelial or vascular smooth muscle cells or both (14) neither predisposes to thrombogenesis nor elevates blood pressure in normolipidemic mice, unlike the same cell selective deletions of COX-2 (15). Global deletion of mPGES-1 reduces atherogenesis and angiotensin II induced aortic aneurysm formation in hyperlipidemic mice, and the proliferative response to vascular injury (14,16,17). This was accompanied by substrate rediversion and augmented formation of PGI_2_, which may contribute to the cardiovascular consequences of mPGES-1 deletion (18–20). In some cell types thromboxane (Tx) A_2_ is the primary rediversion product (16). However, although PGE_2_ is the main prostanoid in inflammatory pain, PGI_2_ has pro-nociceptive actions in some rodent models (21–23) and is the dominant mediator in certain types of human pain (24). Thus, even if substrate rediversion contributes to cardioprotective effects, it might limit the analgesic efficacy of mPGES-1 inhibitors or introduce variability in pain relief.

Here, we engineered mice with deletion of mPGES-1 in myeloid cells, particularly in macrophages as its expression in neutrophils and dendritic cells is low (25), and studied its analgesic and anti-inflammatory effects in models of analgesia. In addition, mice with mPGES-1 conditionally depleted in neurons and glial cells were also generated to determine the relative contribution of peripheral and central mPGES-1 to inflammatory pain hypersensitivity. Our data demonstrate that depletion of peripheral myeloid cell mPGES-1 fully recapitulates the analgesic and anti-inflammatory efficacy of selective COX-2 inhibition. In contrast, central mPGES-1 depletion does not detectably influence inflammatory pain hypersensitivity in mice. This strengthens the therapeutic rationale for targeting myeloid cell, especially macrophage, mPGES-1 in inflammatory pain.

## Methods

### Generation of conditional mPGES-1 deficient mice

As previously described, myeloid cell mPGES-1 deficient mice were generated by crossing mPGES-1flox/flox mice with LysM-Cre mice (26). Selective deletion of mPGES-1 in myeloid cells particularly favors macrophages as it is absent or only trivially present in neutrophils and dendritic cells (25). Moreover, Rittner et al. showed that CFA-induced inflammation, hyperalgesia and PGE_2_ production were independent of infiltrating neutrophils (27). Therefore, the dominant impact in the KOs is on macrophage mPGES-1 (Mac-mPGES-1-KO) (28). LysM-Cre mice without mPGES-1flox/flox were used as controls (Mac-mPGES-1-WT). For neuronal specific mPGES-1 depletion, mPGES-1flox/flox mice were crossed with Nestin-Cre mice to yield mice with mPGES-1 selectively deleted in neurons and glial cells (Neu-mPGES-1-KO) (29). mPGES-1flox/flox mice without Nestin-Cre were used as their controls (Neu-mPGES-1-WT). Mice of age 10-16 weeks were used for all experiments. All procedures were in accordance with the guidelines approved by the University of Pennsylvania Institutional Animal Care and Use Committee.

### Induction of peripheral inflammation

Mice were briefly anesthetized with isoflurane. 1) Paw inflammation was induced by injection of 20 μl of complete Freund’s adjuvant (CFA) (Sigma) subcutaneously into the central plantar region of the left hind paws (29). 2) Tibio-tarsal peri-articular inflammation was triggered by administration of 100μg zymosan A (Sigma) in 20 μl of 0.9% saline into the left tibio-tarsal joint region (30). 3) Collagen II antibody-induced arthritis was achieved as described (31). Briefly, 2 mg monoclonal antibody to collagen II (Chondrex) in 50 ul of PBS was administered by intraperitoneal injection. Three days after the administration of the antibody, mice were injected intraperitoneally with 50 μg of LPS (Sigma) in 0.1 ml of saline to induce the development of arthritis which was primed by the collagen II antibody. Paw and joint thickness were measured using calipers. 4) Formalin induced pain was elicited by intraplantar injection of 15 ul of 5% formalin solution (Sigma) and the time spent licking the injected paw was manually recorded in 5-minute intervals up to 45 minutes after formalin injection. 5) Peritonitis was induced by injection of 1mg of zymosan A in 0.5 ml of saline into the peritoneal cavity.

### Behavioral testing

Mechanical pain hypersensitivity was accomplished with the dynamic plantar von Frey analgesiometer (Stoelting). Mice were placed individually in plastic cages with a wire mesh bottom and allowed to acclimate for 30 minutes. After cessation of exploratory behavior, an automated touch-stimulator (electronic von Frey) was placed below the target area of the inflamed paws and the force increased automatically (linear increase of 0.5 g/s with an upper limit of 5g and a cut off of 20 seconds). Paw withdrawal threshold (PWT) was determined by recording the force at which the animal withdraws its paw. In each test session, each mouse was tested in three sequential trials with an interval of 2-3 minutes.

Thermal pain hypersensitivity was accomplished with a heated glass based plantar analgesiometer (Stoelting). Mice were placed individually in plexiglass cubicles mounted on a glass surface maintained at 30°C and allowed to acclimate for 30 minutes. Then the heat source, a light beam, was focused on the top surface of the glass to create an intense spot of heat under the targeted paw. Paw withdrawal latency (PWL) with a cut off of 20 seconds was defined as the time from onset of stimuli until the mouse withdraws its inflamed paw. Following such a response, the heat source was immediately stopped to prevent tissue damage. In each test session, each mouse was tested in three sequential trials with an interval of 2-3 minutes.

### Wheel running analysis

Wheel running activity was assessed in polycarbonate cages with free access to stainless steel activity wheels which can be turned in either direction as described previously (32). The wheels were connected to a computer that automatically and continuously recorded the rotation counts of each animal in the wheel. Unilateral injection of CFA induced pain did not affect running distances, so bilateral hind paw inflammation was induced for the wheel running experiment.

### DiR-HDL nanoparticle experiments

To assess macrophage accumulation in paw tissues, Mac-mPGES-1-WT and Mac-mPGES-1-KO mice received an i.v. (tail vein) injection of high-density lipoprotein nanoparticles labeled with fluorophore DiR-HDL (D-12731; Invitrogen) in 0.9% sterile saline at 1 μmol DiR/kg as described (25). Immediately after the nanoparticle injection, mice were injected with 20 μl of CFA and then subjected to near infrared fluorescence using an IVIS Spectrum fluorescence imaging system at different time points.

For clodronate liposome-mediated macrophage depletion, Mac-mPGES-1-WT mice were i.v. injected with 200 μl of clodronate liposomes or equal volume of PBS liposomes. 18h later, mice were injected with DiR-HDL nanoparticles as above and 20 ul of CFA was injected immediately after DiR-HDL to induce hind paw inflammation. 24h later, mice were subjected to near infrared fluorescence imaging and then the inflamed soft paw tissue was dissected and OCT-embedded for immunofluorescence staining for macrophage marker CD68.

### Flow cytometry analysis

Paw tissues were minced and digested in 1 mg/ml collagenase A / 2.4 U/ml dispase II (Roche Applied Sciences) in HEPES buffered saline (Sigma) for 2h at 37°C. After digestion, cells were filtered through a 70 μm mesh strainer and centrifuged at 1000 rpm, 5 minutes at 4°C. Following lysis of red cells using 1 ml HBSS (w/2% FBS) and 3 ml NH4Cl, cells were washed and re-suspended in FACS buffer to make a final concentration of 1×10^6^ cells / 100 μl. The suspension was then incubated with antibodies for F4/80 (for macrophages, Invitrogen) and Gr1 (for neutrophils, Invitrogen) after being blocked with an Fc-Blocker (BD Pharmingen). Flow cytometry was performed on BD FACSCalibur and the data was analyzed using the FloJo software (TreeStar Inc.).

Zymosan induced peritoneal cells were collected by using 2 ml of sterile PBS to wash out the inflamed peritoneal cavity. Similarly, after lysis red blood cells, the exudate was re-suspended in FACS buffer and then incubated with F4/80 and Gr1 antibodies for flow cytometry analysis as described above.

### Boyden chamber assay

Thioglycollate-elicited peritoneal macrophages were harvested and cultured as previously described (28). Aliquots of 50,000 macrophages were added to the top wells of Costar Transwell modified Boyden chambers (6.5-mm-diameter tissue culture-treated polycarbonate membranes containing 8-μm pores, Corning). Media with or without 10ng/ml monocyte chemotactic protein-1 (MCP-1) was placed in the lower chambers. After allowing cell migration for 24 hours, cells remaining in the top wells were scraped off using cotton swabs, and all remaining cells were stained with 0.1% crystal violet (Sigma) for 20 minutes. Cells were then washed with 1% deoxycholic acid (Sigma) solution and absorbance was measured at 595nm.

### Mass spectrometry and ELISA analysis

Paws were retrieved and the subcutaneous tissue was cut into small pieces and put in lysis buffer (0.1 M PBS, 10 uM indomethacin, 1 mM EDTA) for mass spectrometry analysis of PGE_2_ and PGI_2_ by liquid chromatography-tandem mass spectrometry (LC-MS/MS) as described (33,34). IL-1β and TNFα were analyzed using ELISA kits (Thermo Scientific) following the manufacturer’s instructions.

### Statistical Analysis

All data are expressed as mean ± SEM or median and interquartile range where appropriate. Statistical differences were determined with two-sample unpaired Student’s t-tests with or without Welch’s correction, depending on whether the variances were significantly different, or with a permutation t-test. P value ≤ 0.05 was considered to be a significant difference.

## Results

### Basal and inflammatory pain hypersensitivity in myeloid cell mPGES-1 deficient mice

To investigate the functional consequences of deletion of mPGES-1 in myeloid cells, we studied mechanical and thermal pain sensitivity in Mac-mPGES-1-KO mice. No significant difference in basal mechanical threshold or thermal latency was detected between Mac-mPGES-1-KO mice and controls (Fig. 1A), indicating minimal contribution of myeloid mPGES-1 to basal nociception. After induction of peripheral hind paw inflammation by intraplantar injection of CFA, the mechanical paw withdrawal threshold (PWT) was markedly reduced in control mice at day 1 and remained decreased over the 21 days of examination, while Mac-mPGES-1-KO mice showed significantly higher thresholds than the control mice across time points (Fig. 1B). Similarly, the paw withdrawal latency (PWL) to thermal stimulation was significantly reduced in control mice, in comparison to Mac-mPGES-1-KO mice (Fig. 1C). CFA injection into the hind paw produces an area of localized peripheral inflammation with persistent paw swelling (increase in paw thickness). Consistent with the reduced pain sensitivity, Mac-mPGES-1-KO mice showed less paw swelling (AUC: 67 ± 2 for KO vs. 76 ± 2 for WT, p=0.0068) (Fig. 1D). These differences were not specific to the CFA-induced hind paw inflammation; similar phenotypes were apparent in two arthritis models, zymosan-evoked peri-articular inflammation (Supplemental Fig. 1A) and collagen II antibody-induced polyarthritis (Supplemental Fig. 1B), which both resulted in significant reductions of mechanical and thermal pain hypersensitivity. Zymosan-evoked joint swelling was also significantly reduced in Mac-mPGES-1-KO mice (Supplemental Fig. 1A).

**Fig 1.**
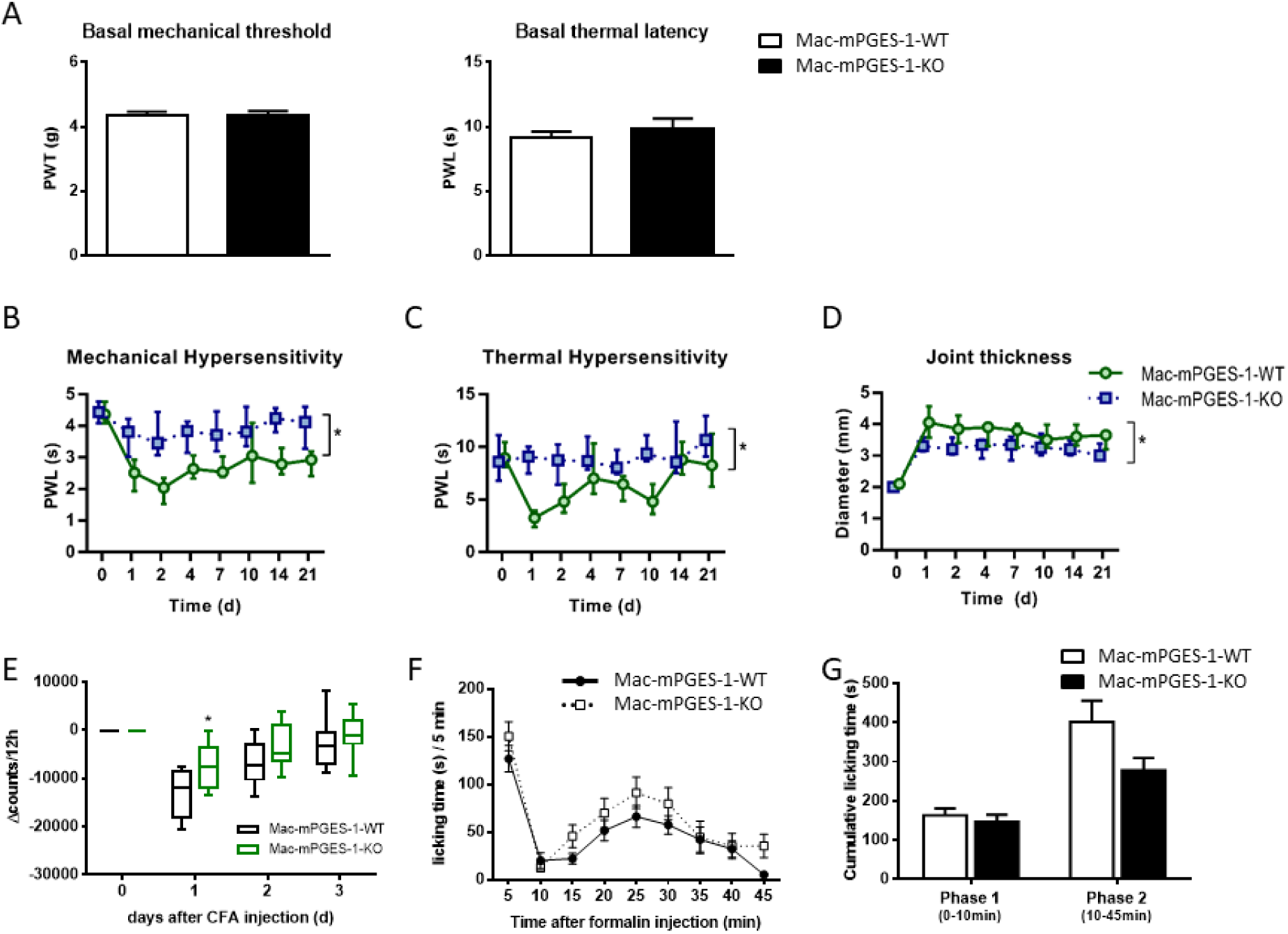
Basal nociception, inflammatory and chemical pain hypersensitivity in Mac-mPGES-1-KO mice. (A), Basal mechanical threshold and thermal latency, n=8. (B-D), Mechanical threshold, thermal latency and paw thickness in CFA-induced hind paw inflammation, n=12 (male & female mice). Median, interquartile range. Statistical test performed on the AUC using a permutation t test. (E), Running wheel activity during the dark phase (7pm-7am) before and after CFA injection, n=9-12. The suppression of activity was reflected by the reduction of rotation counts. (F), Paw licking in 5-minute intervals after intraplantar formalin injection, n=11-12. (G), Cumulative licking time in phase 1 (0-10 minutes) and phase 2 (10-45 minutes) of the formalin reaction, n=11-12.

We next assessed the effect of myeloid mPGES-1 depletion on voluntary wheel running as an index of mobility impairment produced by inflammatory pain (32). The rotation counts over 12 hours (mouse active phase, 7pm-7am) of free access to wheels were significantly decreased in response to CFA-induced hind paw inflammation. Compared to control mice, Mac-mPGES-1-KO mice showed a significant reduction in the decrease in running wheel performance at day 1 (Fig. 1E). These data demonstrate that function, in addition to reflexive measures such as paw retraction, was also significantly influenced by myeloid mPGES-1 depletion.

Formalin-induced chemical pain sensitivity in Mac-mPGES-1-KO mice was similar to control mice (Fig. 1F). Intraplantar injection of formalin evokes 2 phases of spontaneous pain-related behavior, the immediate short-lasting phase (0-10 minutes) due to the activation of peripheral nociceptors, and the second phase (10-45 minutes) which reflects activity-dependent synaptic changes in the CNS (29). We observed no difference in either phase 1 or phase 2 between Mac-mPGES-1-KO and control mice (Fig. 1G), implying a limited contribution of myeloid cell mPGES-1 to activity-dependent central sensitization.

### Basal and inflammatory pain hypersensitivity in neuronal mPGES-1 deficient mice

Neuronal specific mPGES-1 depletion (Neu-mPGES-1-KO) mice were used to study the role of central mPGES-1 in inflammatory pain hypersensitivity. mPGES-1 expression was significantly inhibited in spinal cord but not in paw tissues in Neu-mPGES-1-KO mice (Fig. 2A), while COX-2 expression was not affected in either tissue (Fig. 2B). Basal and inflammatory pain responses were assessed in these mice. No difference in basal mechanical or thermal threshold was detected between Neu-mPGES-1-KO and control mice (Fig. 2C). In contrast to Mac-mPGES-1-KOs, the Neu-mPGES-1-KO mice did not display any difference in mechanical threshold (Fig. 2D), thermal latency (Fig. 2E) or paw thickness (Fig. 2F) in CFA-induced hind paw inflammation. No detectable phenotypic impact on peripheral inflammation and the consequent hyperalgesia was observed in collagen II antibody-induced polyarthritis models as well (Supplemental Fig. 2).

**Fig 2.**
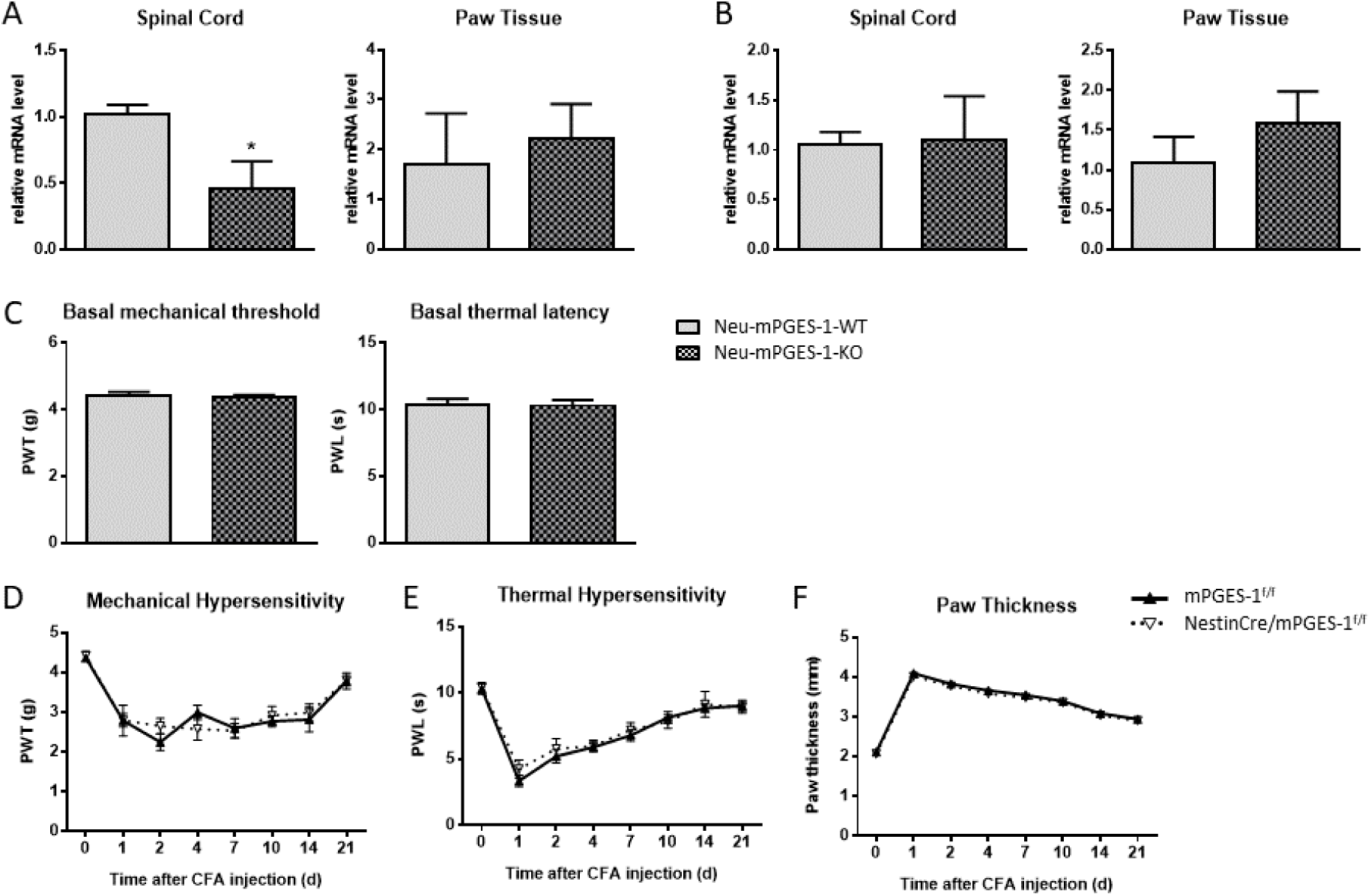
Basal nociception and inflammatory pain hypersensitivity in Neu-mPGES-1-KO mice. (A-B), Real-time PCR analysis of mPGES-1 (A) and COX-2 (B) mRNA expression in spinal cord (n=5-7) and paw tissues (n=3-4). (C), Basal mechanical threshold and thermal latency, n=9. (D-F), Mechanical threshold, thermal latency and paw thickness in CFA-induced hind paw inflammation, n=9.

### Inhibition of COX-2 does not reduce inflammation and hyperalgesia more effectively than myeloid mPGES-1 depletion

To assess if myeloid cell mPGES-1 depletion would recapitulate the analgesic efficacy of selective COX-2 inhibition, or if blockade of COX-2 could further improve inflammatory hyperalgesia in Mac-mPGES-1-KO mice, we fed Mac-mPGES-1-WT and KO mice with celecoxib diet at a biochemically-validated COX-2 selective concentration (100 mg/kg/day). After 3 weeks of celecoxib diet, mice were injected with CFA to induce hind paw inflammation. Mechanical and thermal pain sensitivity was then assessed for 7 consecutive days. In Mac-mPGES-1-WT group, celecoxib treatment significantly increased the mechanical pain threshold (Fig. 3A, p<0.001), prolonged the thermal pain latency (Fig. 3B, p<0.05), and attenuated peripheral paw swelling (Fig. 3C, p<0.01). By contrast, celecoxib did not further improve CFA-induced paw swelling and pain thresholds in Mac-mPGES-1-KO mice (Fig. 3A-C, p>0.05 KO celecoxib vs. KO ctrl); both groups displayed effects equivalent to the celecoxib treatment in WT mice (Fig. 3A-C, p>0.05 KO celecoxib vs. WT celecoxib; p>0.05 vs. KO ctrl vs. WT celecoxib).

**Fig 3.**
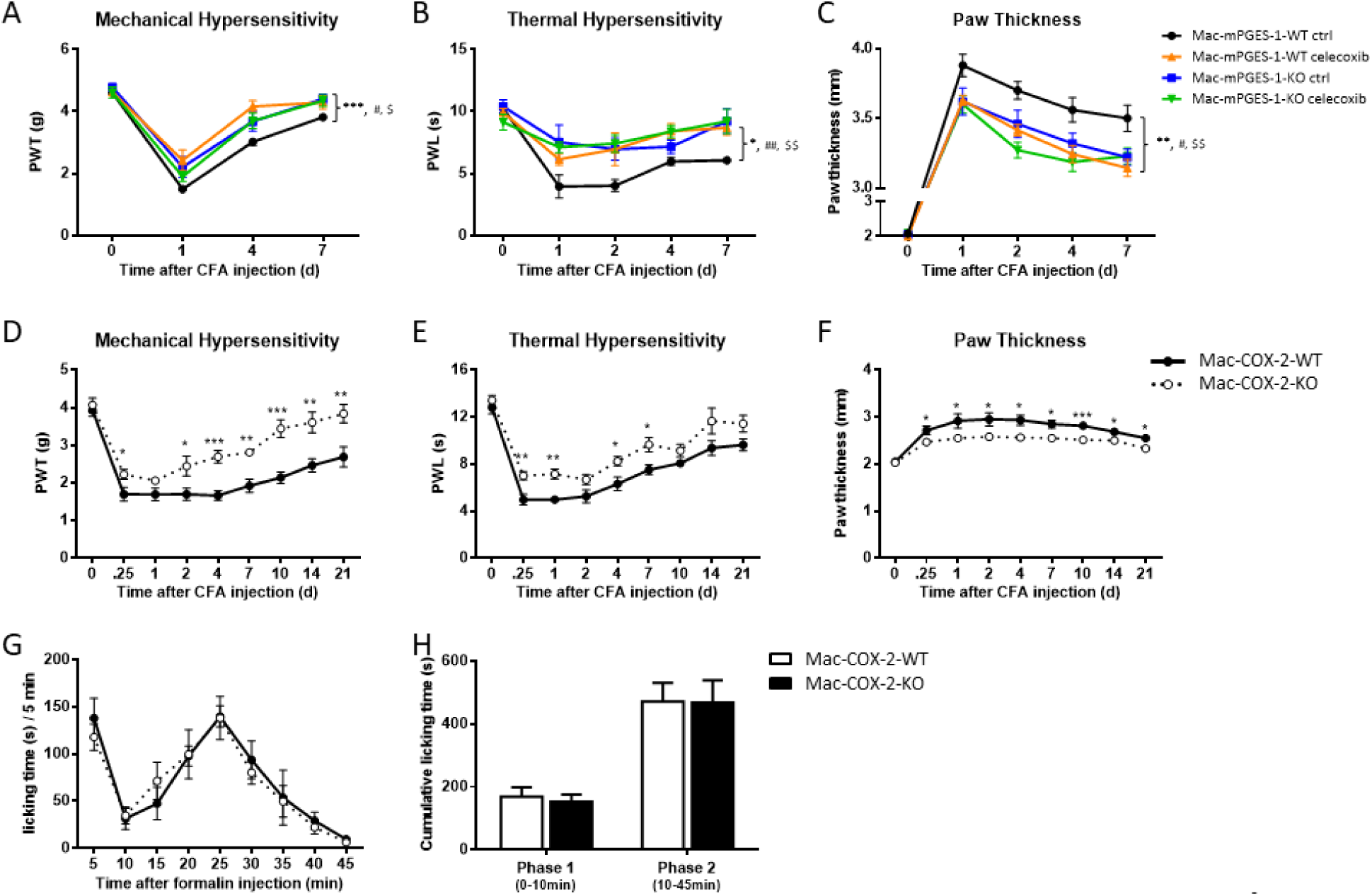
Blockade of COX-2 has no further effects on the development of peripheral inflammation and pain hypersensitivity in Mac-mPGES-1-KO mice. (A-C), Mac-mPGES-1-KO and Mac-mPGES-1-WT mice were fed with celecoxib diet (100mg/kg/day) for 3 weeks before the induction of CFA-induced hind paw inflammation. Mechanical threshold, thermal latency and paw thickness before and after CFA injection, n=5-7. (D-F), Mechanical threshold, thermal latency and paw thickness after CFA-induced hind paw inflammation in Mac-COX-2-KO and control mice, n=6. (G), Paw licking in 5-minute intervals after intraplantar formalin injection in Mac-COX-2-KO and control mice, n=5. (H), Cumulative licking time in phase 1 (0-10 minutes) and phase 2 (10-45 minutes) of the formalin reaction in Mac-COX-2-KO and control mice, n=5.

Furthermore, we measured the inflammation and pain responses in mice which lack COX-2 in myeloid cells, particularly in macrophages (Mac-COX-2-KO) (28). As shown in Figure 3D-F, the phenotypes we observed in Mac-COX-2-KO mice recapitulated those seen in Mac-mPGES-1-KO mice with CFA-induced hind paw inflammation, including increased threshold to mechanical pain stimuli (Fig. 3D), prolonged latency to thermal pain stimuli (Fig. 3E) and less peripheral paw swelling (Fig. 3F). Similarly, pain behavior evoked by intraplantar formalin injection did not reveal obvious differences between the Mac-COX-2-KO and controls (Fig. 3G), neither in first nor second phase responses (Fig. 3H).

### Myeloid mPGES-1 deficiency suppresses the production of PGE_2_ and inflammatory cytokines

To study the underlying mechanisms of the reduced pain and inflammation in myeloid mPGES-1 KO mice, we measured the production of PGE_2_ in mouse paw tissues. CFA stimulation evoked marked PGE_2_ production in control paws after 24 hours, while this increase was reduced in the Mac-mPGES-1-KO paws (Fig. 4A). CFA-evoked secretion of pro-inflammatory cytokines, such as IL-1β (Fig. 4B) and TNF-α (Fig. 4C), was also significantly inhibited in Mac-mPGES-1-KO paws.

**Fig 4.**
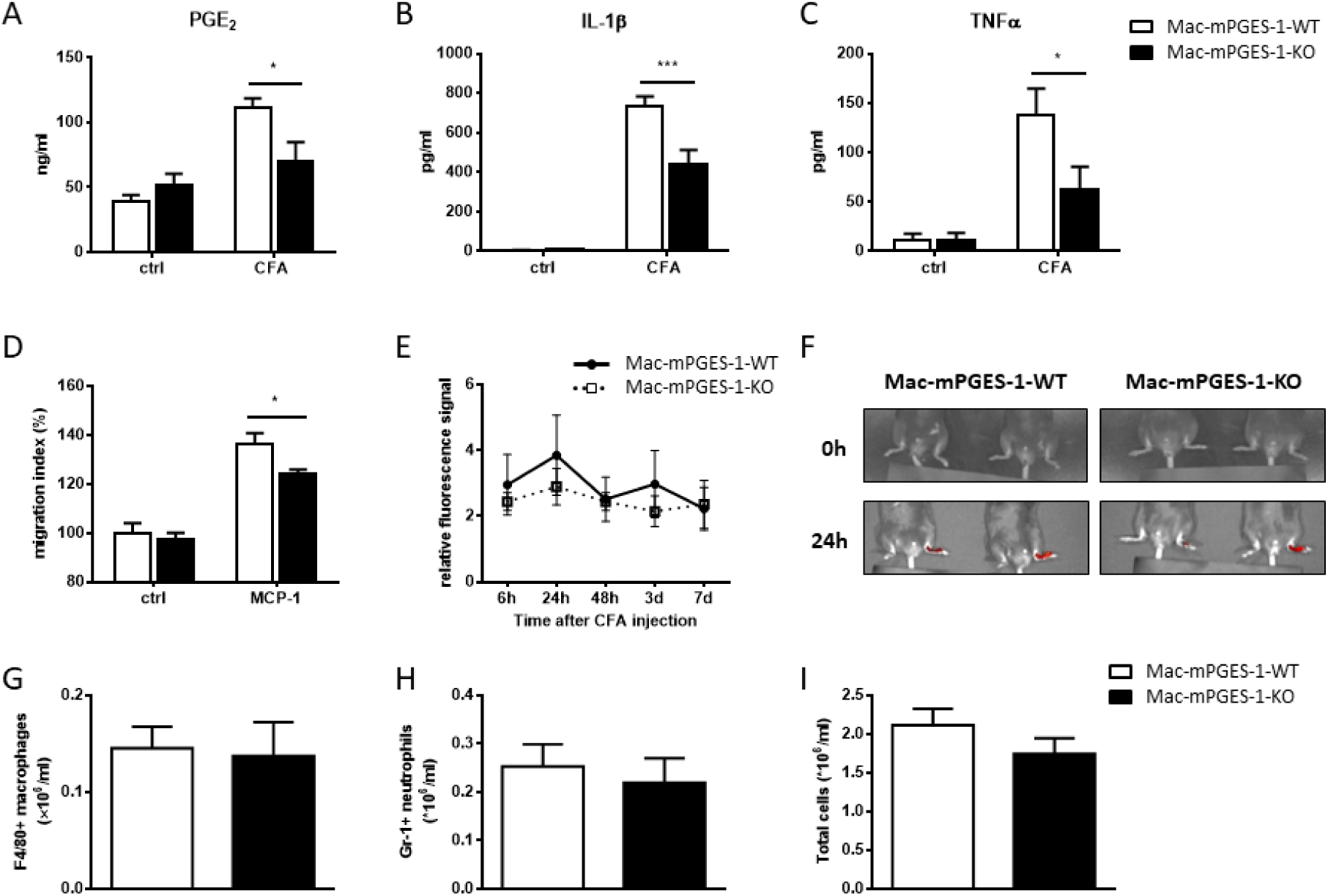
(A-C), PGE_2_, IL-1β and TNFα levels in paw tissues before and 24h after CFA injection in Mac-mPGES-1-KO mice, n=4-8. (D), Deficiency of mPGES-1 impairs macrophage migration potential in vitro. Migration index of the WT cells without MCP-1 is defined as 100%, n=17. (E), Time course of the fluorescence signal in paws from mice injected with CFA and DiR-HDL nanoparticles, n=6-7. Data were calculated using the formula: (SI(lp)-SI(rp))/(SI(rp)-SI(rp-preinjection)). SI: signal intensity; lp: left inflamed paw; rp: right uninjured paw. (F), Representative images of paws from mice injected with DiR-HDL nanoparticles before and 24h after CFA injection. (G-I), Flow cytometry analysis of the amounts of F4/80+ macrophages, Gr1+ neutrophils and the total infiltrated cells in inflamed paw tissues 48h after CFA injection, n=12.

### Effect of myeloid mPGES-1 deletion on macrophage accumulation

Inflammatory cell recruitment is a hallmark of the development of inflammation. Activation of macrophages by inflammatory stimuli markedly promotes the expression of COX-2 and mPGES-1 and augments the subsequent release of its major product, PGE_2_, which mediates pain and inflammation (35). To evaluate if the reduced PGE_2_ and cytokine production are attributable to reduced macrophage accumulation at the inflamed sites, we first performed an in vitro Boyden chamber assay to detect the effect of mPGES-1 depletion on macrophage migration potential. As shown in Fig. 4D, mPGES-1 deficiency did not change random macrophage migration at baseline, while MCP-1-directed macrophage migration was significantly inhibited by 11.7% ± 0.5% (p<0.05) in Mac-mPGES-1-KO cells. However, the macrophage content paw tissue in vivo was not significantly reduced in the KOs (Fig. 4E & F) as determined by fluorescence imaging of the distribution of 1,1-dioctadecyl-3,3,3,3-tetramethylindotricarbocyanine iodide (DiR-HDL) nanoparticles. DiR-HDL nanoparticles accumulated preferentially in the inflamed paws while the uninjured control paws showed no signal above background. The specific targeting to macrophages of these DiR-HDL nanoparticles was confirmed by the co-localization of DiR and the macrophage marker CD68 (Supplemental Fig. 3A) and by the significantly reduced fluorescence signal when macrophages were systematically depleted by treatment with clodronate liposomes (Supplemental Fig. 3B).

Likewise, ex vivo flow cytometry failed to detect differences in macrophage infiltration between WT and Mac-mPGES-1-KO paws 48h after CFA treatment (Fig. 4G). Neutrophil content (Fig. 4H) and the total infiltrated cells (Fig. 4I) also showed no difference. We further independently assessed the peritoneal inflammatory cell infiltration in a zymosan-induced peritonitis model. Although the neutrophil content showed a reduction 24h after peritonitis induction in Mac-mPGES-1-KO mice, the macrophage content was not different either at 24h or 48h (Supplemental Fig. 4). These data collectively demonstrate that the macrophage recruitment was unaltered by myeloid mPGES-1 depletion in vivo, and that the impact on pain and inflammation in Mac-mPGES-1-KO mice may reflect a change in macrophage functions, such as reduced production of PGE_2_ and inflammatory cytokines, rather than in number.

## Discussion

Prostaglandin E2 (PGE_2_) plays an important role in regulating the sensitivity of sensory neurons and enhancing pain perception during inflammation (36). The efficacy of drugs targeting the PGE_2_ pathway by systemic or intrathecal administration of traditional NSAIDs or COX-2 selective inhibitors, is well established in inflammatory pain (37–39). However, due to the concurrent inhibition of cardioprotective prostacyclin (PGI_2_), patients taking COX-2 inhibitors, such as celecoxib or older drugs like diclofenac, are predisposed to a cardiovascular hazard, including hypertension, stroke, myocardial infarction, heart failure and sudden cardiac death (40,41). Therefore, interest has focused on mPGES-1 as an alternative target for the development of analgesic and anti-inflammatory drugs. Indeed, global deletion of mPGES-1 has effects similar to NSAIDs in rodent models, such as acetate-elicited acute pain responses (8,9), and collagen- or collagen antibody-induced chronic arthritis (42,43).

In the current study, we examined the specific role of myeloid cell mPGES-1, particularly in macrophages, in inflammatory pain. Compared to controls, absence of mPGES-1 in myeloid cells resulted in significant relief of both mechanical and thermal pain hypersensitivity in both soft tissue inflammation and arthritis models. Peripheral inflammation reflected by local tissue swelling was also significantly reduced in Mac-mPGES-1-KO mice. These effects were reflected by changes in macrophage functions, but not macrophage recruitment. Thus, concomitant with the suppression of PGE_2_ production and inflammatory cytokine secretion, peripheral macrophage accumulation was unaltered in KOs. Although there was a marked reduction of pain hypersensitivity in Mac-mPGES-1-KO mice after induction of peripheral inflammation, we found no difference between control and KO mice at baseline in either mechanical or thermal pain sensitivity, indicating minimal contribution of myeloid mPGES-1 to basal nociception. Also, these mice showed comparable formalin-induced chemical pain sensitivity to control mice. Formalin injection usually evokes 2 phases of spontaneous pain-related behavior (29,44), an immediate short-lasting first phase due to the direct activation of TRPA1-expressing nociceptors and a slower-onset, longer-lasting phase 2 that reflects central sensitization in the spinal cord. These results imply a limited contribution of constitutive myeloid mPGES-1 to activity-dependent central sensitization. Similarly, myeloid COX-2 deletion did not alter formalin-induced pain sensitivity. Considering the same minor contribution of central COX-2 to formalin-mediated pain behavior (29), the response to the formalin test might rely mainly on constitutive COX-1 (45,46), although others point to a role for COX-2 (47) or an involvement of both (48).

We compared the analgesic effects of myeloid mPGES-1 deficiency with the impact of a COX-2 selective inhibitor, celecoxib. Celecoxib is the only NSAID developed purposefully to inhibit selectively COX-2 that remains on the US market, and it attenuates pain and inflammation in mice. Here we showed that celecoxib failed to further improve CFA-induced inflammation (paw thickness) and hyperalgesia (mechanical and thermal pain hypersensitivity) in Mac-mPGES-1-KO mice. These data support that myeloid mPGES-1 derived PGE_2_ is the functionally dominant prostanoid in the inflammatory pain models studied here, because COX-2 inhibition in macrophages also inhibits the production of other prostanoids (28), especially PGI_2_, which is also considered as an important pain mediator (21). In contrast, lack of mPGES-1 in macrophages selectively inhibits PGE_2_ synthesis, but is accompanied by an increase in PGI_2_ likely through a mechanism of substrate rediversion in the PGI_2_ biosynthetic pathway (49). Thus, the observed similar analgesic effects of myeloid COX-2 and mPGES-1 deficiency demonstrate that PGE_2_, but not PGI_2_, in myeloid cells is the major mediator of inflammatory pain. Suppression of the myeloid mPGES-1-PGE_2_ pathway may account for most, if not all, of the analgesic efficacy of COX-2 blockade. This supports that selectively targeting myeloid mPGES-1 might preserve the analgesic efficacy of NSAIDs.

Growing evidence indicates that the effect of PGE_2_ in enhancing pain perception during inflammation occurs both at the peripheral site of injury and in the spinal cord (50–53). We excluded the contribution of central mPGES-1 to inflammatory pain by using mice with mPGES-1 selectively deleted in neurons and glial cells, further demonstrating that peripheral macrophage mPGES-1 derived PGE_2_ is the dominant source of prostanoids in inflammatory pain. In contrast to the null phenotype of neuronal mPGES-1 deficiency, Vardeh et al. demonstrated a pivotal contribution of neuronal COX-2 to inflammatory mechanical pain hypersensitivity (29). Other prostanoids redirected from PGE_2_ in neuronal mPGES-1 null mice which also serve as nociceptive mediators (54,55) and maintain the pain response might contribute to this apparent discrepancy. The different expression profiles (56,57) and diverse biological activities (58–61) of the 4 PGE_2_ receptors in the spinal cord i might be another potential explanation. Furthermore, although the neuronal COX-2 deficient mice showed complete loss of mechanical hypersensitivity in a peripheral inflammatory pain model (29), our data revealed that myeloid cell COX-2/mPGES-1 also contribute to inflammatory mechanical pain. Similarly, although the role of neuronal COX-2 in thermal pain is being debated (29,62), our results support an important role of myeloid COX-2/mPGES-1 in the response to thermal stimuli.

Notably, the relief of pain and reduction of inflammation by myeloid COX-2 or mPGES-1 depletion was not complete in our settings; the overall peripheral inflammation and pain still remained detectable, implying that myeloid COX-2-mPGES-1-PGE_2_ pathway is not the sole contributor. COX-2/mPGES-1 expression in other immune cells, e.g. T cells and B cells, or in injured endothelial cells, might also play important roles; selective deletion of COX-2/mPGES-1 in these cells will be required for further studies. Moreover, although the expression of COX-2/mPGES-1 in these cells appears variable (45,63–66), we cannot exclude the contribution of COX-2/mPGES-1 in microglia, the resident macrophages in the CNS. Future experiments utilizing a more specific microglia-Cre driver will be needed to determine the precise anatomical locus of the effect.

In summary, we show that the myeloid COX-2-mPGES-1-PGE_2_ biosynthetic pathway plays an essential role in the development of peripheral inflammation and related pain hypersensitivity in mouse models. Considering the relatively benign cardiovascular profile of myeloid mPGES-1 depletion in mice, our observations strengthen the rationale for targeting myeloid cell mPGES-1.

## Acknowledgements

We thank Dr. Mohammad Bohlooly and Dr. Xiufeng Xu from Astra Zeneca for kindly providing the mPGES-1flox/flox mice. We thank John Lawson, Helen Zou and Wenxuan Li for their technical support with the mass spectrometry analysis and thank Sarah Teegarden for her assistance with the manuscript writing. This work was supported by the National Heart, Lung, and the Blood Institute funded Personalized NSAID Therapeutics Consortium (HL117798; T.G., G.A.F.).

