## Supplemental Fig for "Myeloid cell mPGES-1 mediates inflammatory pain hypersensitivity in mice"

**Supplemental Figures**

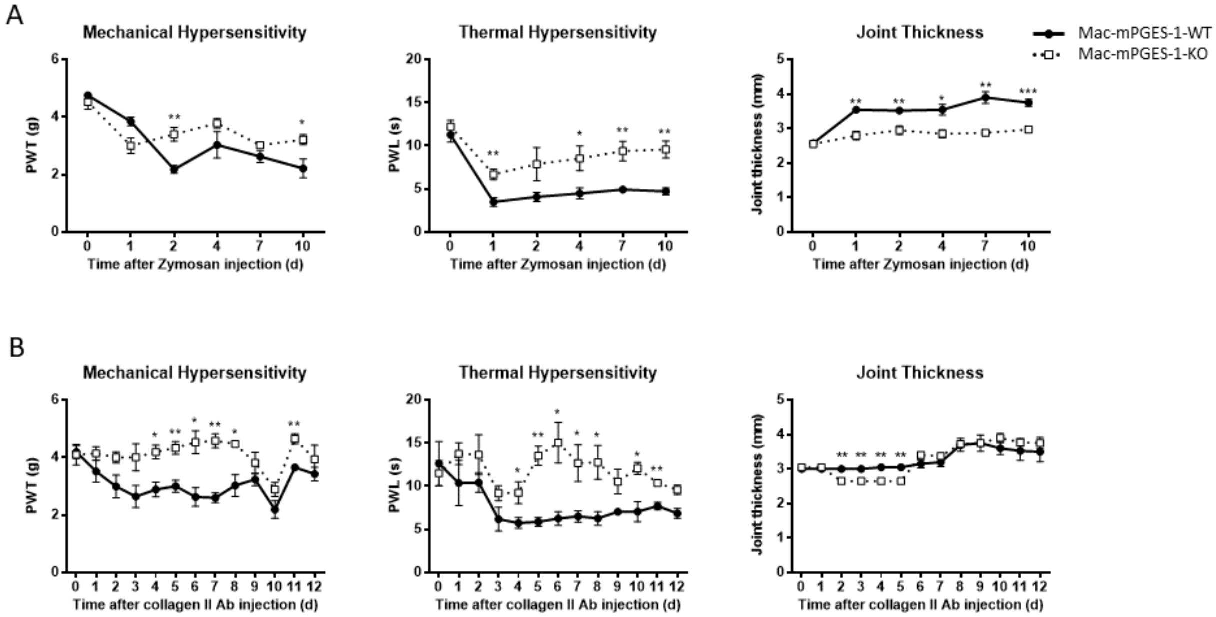


Supplemental Fig. 1. Mechanical threshold, thermal latency and tibiotarsal joint thickness in Mac-mPGES-1-KO mice with zymosan-induced peri-articular inflammation (A) and collagen II antibody-induced polyarthritis (B), n=4.

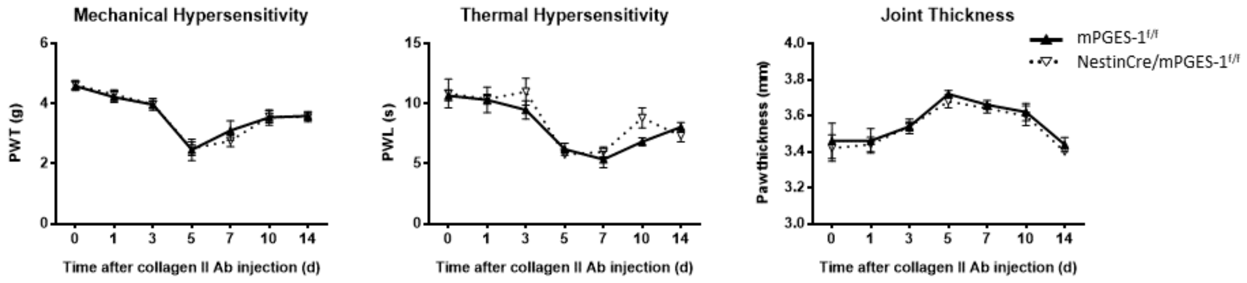


Supplemental Fig. 2. Mechanical threshold, thermal latency and tibiotarsal joint thickness in Neu-mPGES-1-KO mice with collagen II antibody-induced polyarthritis, n=5.

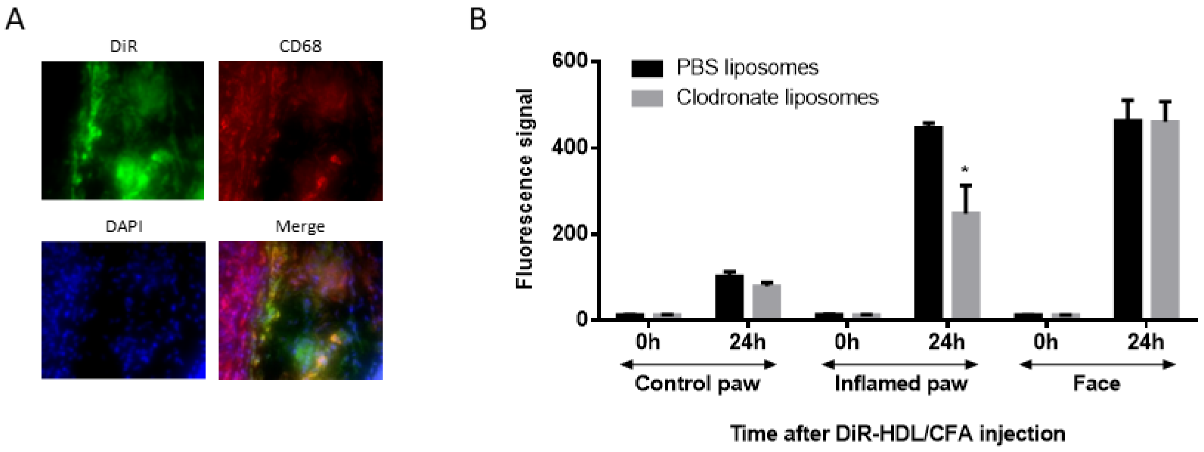


Supplemental Fig. 3. (A), Immunofluorescence validated the co-localization (Merge, yellow) of the fluorescent dye DiR (green) with the macrophage marker CD68 (red) in inflamed paw sections. DAPI (blue) indicated nuclear staining. (B), 24h after CFA injection, DiR-HDL fluorescence signal was significantly reduced in inflamed paws from mice with clodronate liposomes injections, while the signal in uninjured control paws and faces was not affected by clodronate liposomes treatment, n=3.

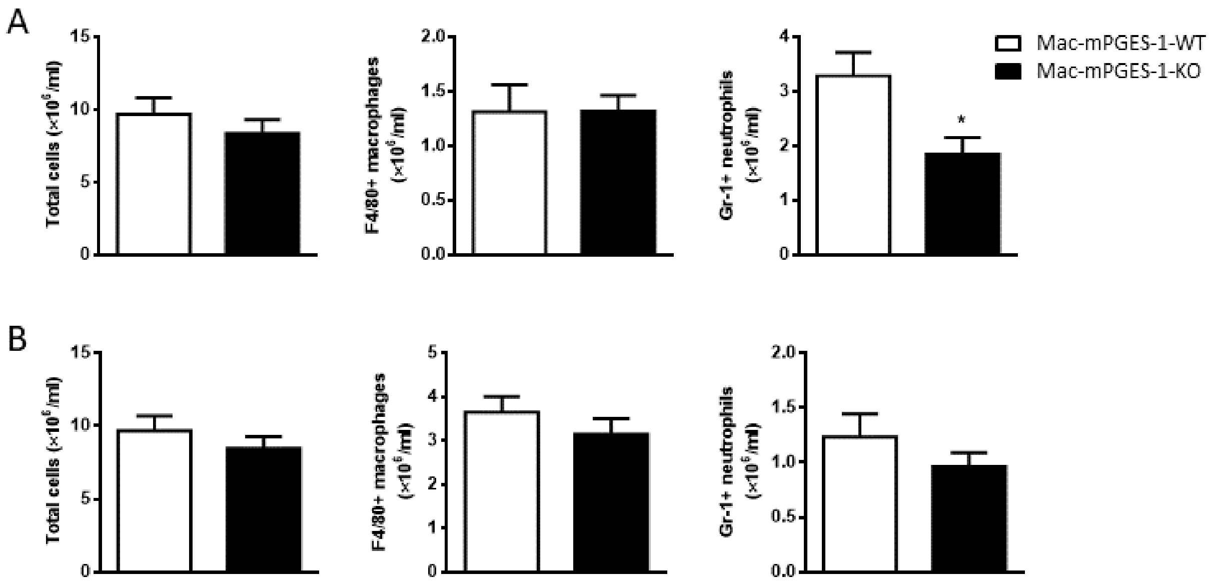


Supplemental Fig. 4. Flow cytometry analysis of the amounts of total infiltrated cells, F4/80+ macrophages and Gr1+ neutrophils in the peritoneal cavity 24h (A) or 48h (B) after zymosan induction of peritonitis, n=16-17.
